# Leptin promotes social context-specific increase in advertisement song effort of male Alston’s singing mice

**DOI:** 10.1101/2025.08.25.672212

**Authors:** Joel A. Tripp, Keerthana Raghuraman, Ritika V. Bhalla, Steven M. Phelps

## Abstract

Animal display behaviors, such as advertisement songs, are flashy and attention grabbing by necessity. In order to balance the costs and benefits of such signals, individuals must be able to assess both their own energetic state and their social environment. In this study, we investigated the role of leptin, a hormonal signal of high energy balance, in regulating the vocal advertisement display of Alston’s singing mouse (*Scotinomys teguina*). We manipulated perception of energy balance using exogenous leptin, and social environment through acoustic playback, to ask how internal cues of energy availability are integrated with external social cues that promote singing. We found that both song playback and leptin injection promoted increased song effort. In the absence of song playback, leptin altered amplitude modulation in songs, but did not affect song rate. Additionally, we examined hormone and playback effects on non-vocal behaviors and found that leptin may shift physical activity away from cage exploration and toward wheel running. Finally, we found several positive associations between measures related to high song effort. These results demonstrate that male singing mice use both social context and energy balance to govern their investment in advertisement song and that leptin acts as a mediator of this process.

## Introduction

Vocal advertisement displays are often critical for resource defense and reproductive success. While the benefits of these behaviors are clear, they also come with inherent costs. Aside from the metabolic expenses of producing a vocalization, which vary by species (Collier et al., 2022; Hasselquist and Bensch, 2008; Ilany et al., 2013; Prestwich, 1994; Ward et al., 2004), these displays can also attract unwanted attention. For example, Neotropical fringe-lipped bats (*Trachops cirrhosus)* locate palatable frog prey using the frog’s mating calls (Tuttle and Ryan, 1981) while the songs of crickets attract parasitoid flies seeking hosts for their larvae (Cade, 1975). Within species, displays may trigger aggression, such as the territorial disputes of calling bullfrogs (Emlen, 1968) or singing birds (Illes and Yunes-Jimenez, 2009; Koloff and Mennill, 2011; Mennill, 2006). Further, engaging in displays requires callers to forego opportunities to forage and increase their energy reserves. Thus, individuals require knowledge of their environment to determine whether an intended receiver is likely to hear their display while also assessing their internal state, including available energy reserves, in order to determine whether they are sufficiently prepared to deal with potential negative consequences of vocalizing.

One important signal of energy balance is the hormone leptin. Leptin is released by fat cells and signals high energy availability (Friedman, 2019). Injection of leptin results in decreased feeding, increased energy expenditure, and reduced body weight (Döring et al., 1998; Halaas et al., 1995; Pelleymounter et al., 1995). While leptin was originally investigated primarily for its role in regulating feeding and its relationship to obesity, it has since also been found to influence puberty onset and fertility (Childs et al., 2021), the stress response (Roubos et al., 2012), and immune function (Abella et al., 2017). Manipulations of circulating leptin have also been shown to influence social and communication signals in electric fish (Sinnett and Markham, 2015) and vocal advertisement displays in rodents (Burkhard et al., 2018; Giglio and Phelps, 2020).

In this study, we investigated the role of leptin and its integration with social context in regulating the vocal display of Alston’s singing mice (*Scotinomys teguina*). Singing mice are small rodents native to Central America, named for their distinctive vocalization (Fig. 1A,B), which consists of a trill of notes that sweep from the ultrasonic to human-audible range. Males increase their rate of singing in response to cues that females are nearby and in response to playback of conspecific song (Fernández-Vargas et al., 2011; Pasch et al., 2017; Tripp and Phelps, 2025), suggesting that as in advertisement displays of other species, song plays a role in both mate attraction and intra-sexual competition. Additionally, high leptin levels have previously been associated with high song effort in singing mice (Burkhard et al., 2018) and exogenous leptin increases the rate of singing (Giglio and Phelps, 2020). These features make the singing mouse an ideal model to investigate the integration of social environment with signals of internal state to regulate social display.

**Fig. 1.**
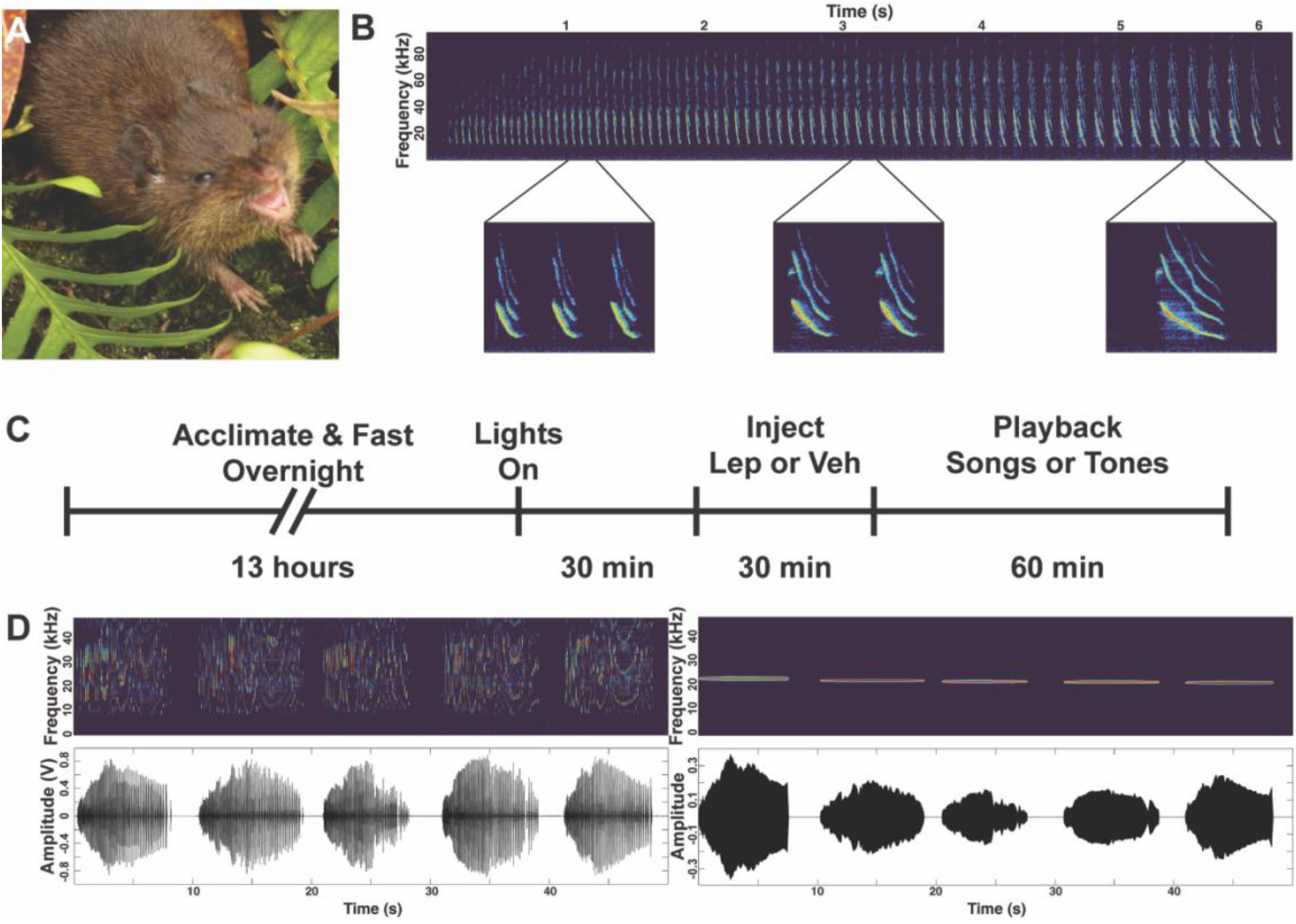
Experiment subject and design. **A)** Singing mouse (*Scotinomys teguina*). **B)** Singing mouse song. Examples of notes at the beginning, middle, and end of song are shown in boxes below. **C)** Experiment timeline. **D)** Spectrograms (above) and oscillograms (below) of example stimulus sets. Set of five stimulus songs on the left and matched control tones on the right.

We conducted an experiment using hormonal manipulations and acoustic playback to better understand how leptin is integrated with external cues of social context to regulate acoustic display investment. In our 2 x 2 design, individuals were first fasted overnight, then received injection of either leptin or vehicle, followed by playback of conspecific songs or control tones. We then examined the previously-identified connections between leptin and song playback on song rate and length (Giglio and Phelps, 2020) and explored the effect of our manipulations on additional song features previously associated with high song effort in wild singing mice (Burkhard et al., 2023, 2018) as well as non-vocal behaviors related to energy use, exploration, and consumption. Our findings deepen our understanding of the role of leptin in social communication and raise new questions about how individuals integrate signals of their own internal state with cues from the environment to regulate social behaviors.

## Material and Methods

### Animal subjects

Subjects were adult (6-32 weeks of age) male singing mice bred in a captive colony derived from wild caught animals collected in the San Gerardo de Dota Valley, Costa Rica. Prior to the experiment, animals were single- or pair-housed in acrylic hamster cages with running wheel and rodent igloo, PVC pipe, and cotton nestlets for enrichment with a 12h:12h light schedule. Pair-housed animals were separated before the beginning of the experiment. Because singing mice are native to montane cloud forests, colony and procedure rooms were kept at 20°C and a small ball of sphagnum moss was included in cages and regularly misted to maintain humidity. All procedures followed the National Institutes of Health Guide for the Care and Use of Laboratory Animals and were approved by the Institutional Animal Care and Use Committee of the University of Texas at Austin.

### Hormone manipulation and acoustic playback

In order to maximize the potential effects of leptin on song effort, each animal was fasted overnight followed by hormone manipulation and acoustic playback in a 2 x 2 design. Fourteen hours prior to playback, we transferred subject animals to a clean home cage without food and isolated them in an acoustic chamber overnight. All other standard enrichment items (see above) and *ad lib* water were provided. The following morning, we opened the acoustic chamber door at the onset of the light period (0900h) so that subject animals could hear playback stimuli (see below) and that we could record video of the subject’s behavior (Fig 1C). At 0930h animals received an injection of either leptin (Peprotech, Cranbury, NJ) in 0.9% saline+0.1% bovine serum albumin vehicle (Sigma-Aldrich, St. Louis, MO) at 10mg/kg or an equivalent volume of vehicle alone.

Playback procedures were adapted from previous studies (Okobi et al., 2019; Tripp and Phelps, 2024). One hour of playback began at 1000h. Playback stimuli consisted of either songs recorded from wild-caught animals unfamiliar to subjects, or control tones which were generated to match the length, mean frequency, and amplitude envelope of stimulus songs (Fig. 1D). Stimuli were played from a speaker in a neighboring acoustic chamber, which elicits singing from subjects more strongly than playback from a nearby speaker (Fujishima and Long, 2025). Each stimulus set contained either five songs recorded from a single animal or five corresponding tones. Acoustic stimuli were played a total of ten times at randomized intervals over the course of the playback hour. We used stimulus sets recorded from four stimulus animals, with the same song and tone sets used within each block of the experiment. The order of hormone treatment and playback stimulus was counterbalanced across blocks.

We recorded vocal behavior using a microphone and preamplifier (ACO Pacific) connected to a Tucker-Davis MA3 amplifier and RX8 microprocessor at 32-bit resolution and sampling rate of 97.7 kHz. Playback and audio recording were controlled by custom Matlab (v9.7.0) and RPvdsEx (v92) scripts. Additional behaviors were recorded using a video camera (Sony Handycam DCR-SR68).

### Song analysis

Following previous methods (Tripp and Phelps, 2025, 2024), we used custom Matlab scripts to bandpass filter recording files at 5-48kHz, then square and down sample to 1000Hz using *resample*. Songs were identified as samples passing an empirically determined threshold and lasting at least three seconds. We categorized vocalizations with a gap of one-half second or longer as separate songs. Measures of song effort, spectral features, and frequency modulation were extracted as previously described (Burkhard et al., 2018; Campbell et al., 2010). Many song features, such as note length, note duration, and inter-note interval change throughout the song (Fig. 1B) and can be modeled using quadratic functions with the “c” term describing the starting value, the “b” term describing the starting slope, and the “a” term describing the rate of change of the slope from note to note.

In this study, we focused on measures associated with high song effort in wild-caught singing mice (Burkhard et al., 2023, 2018), specifically the number of songs during playback, song length, number of notes per song, starting inter-note interval (INIc), and two measures of amplitude modulation: starting slope of relative peak amplitude (PAb), and rate of change of relative peak amplitude slope (PAa). We chose to focus on relative amplitude measures because the distance and orientation of the mouse can vary, making absolute amplitude measures unreliable. To account for this, we normalized the peak amplitude of each recording to a value of 1 (Campbell et al., 2010). In addition to these measures previously associated with song effort, we also examined the rate and latency of counter-singing, which was defined as the number of times a playback was followed by a song within 20 s of playback onset (Tripp and Phelps, 2024). One recording file was truncated and could not be processed using these methods. For that subject, in the leptin and song playback group, we identified number and duration of songs using video recordings and excluded it from all other analyses.

### Non-vocal behavior analysis

We used BORIS (v. 7.8.2; Friard and Gamba, 2015) to confirm behavioral analyses from audio recordings and quantify non-vocal behaviors including time spent running on the exercise wheel, exploring the cage, defined as time actively moving around the cage but not running on the wheel, and drinking water. Videos were scored by two observers blind to subject condition. To characterize inter-rater reliability for each behavior of interest, we calculated Pearson’s correlations for total duration quantified by the two observers and found a significant positive correlation for each (Wheel running: R = 0.99, P = 1.24×10^-9^; Exploring: R = 0.64, P = 0.024, Drinking: R = 0.98, P = 1.15×10^-8^). These results indicate a high degree of agreement for wheel running and drinking and weaker agreement for exploring. For blocks analyzed by both observers, we used the average duration value for each behavior.

### Statistical analysis

We used the RStudio environment (v 2023.06.0+421) for R (v4.2.2) to conduct statistical analyses and to generate figures. Song counts were fit to a generalized linear model with a Poisson distribution using the *lme4* package (Bates et al., 2015). We tested for differences across hormone condition, playback condition, and a hormone x playback interaction using an ANOVA with type III sum of squares implemented by the *car* package (Fox et al., 2024). Pairwise comparisons across groups were made using the *emmeans* package (Lenth et al., 2022) with a significance cut-off of 0.05 for Tukey-adjusted p-values. Number of counter-songs produced during the playback hour were compared for animals receiving song playback using generalized linear model with quasipoisson distribution. Counter-song latencies were compared using a two-sided t-test. For song structure measures (song length, notes per song, INIc, PAb, and PAa) we compared the mean value from each subject, excluding subjects that did not sing during the playback hour. These measures, along with non-vocal behaviors (time running on wheel, time exploring cage, and time drinking water) were each compared using two-way ANOVA and TukeyHSD was used for post hoc comparisons. We calculated pairwise Spearman correlations between vocal and non-vocal behavioral measures using the *psych* package (Revelle, 2025) using the Benjamini-Hochberg method for False Discovery Rate correction (Benjamini and Hochberg, 1995). As with song structure measures, we excluded mice that did not sing from correlational analyses.

## Results

### Leptin increases singing rate in response to song playback

Both leptin treatment and song playback increased the rate of singing in male mice (Fig. 2A). ANOVA applied to our generalized linear model showed a significant effect of hormone condition (F_1,44_ = 4.95, p = 0.031) and playback condition (F_1,44_ = 16.32, p = 0.00021), but no interaction effect (F_1,44_ = 0.15, p = 0.70). *Post hoc* tests revealed that animals treated with leptin and given song playback sang significantly more than all other groups (p < 0.0001 for all comparisons), while animals receiving vehicle injection and song playback sang significantly more than either tone playback group (p < 0.0001 for both comparisons), and animals that received tone playback sang at similarly low rates, regardless of hormone treatment (p = 0.64).

**Fig. 2.**
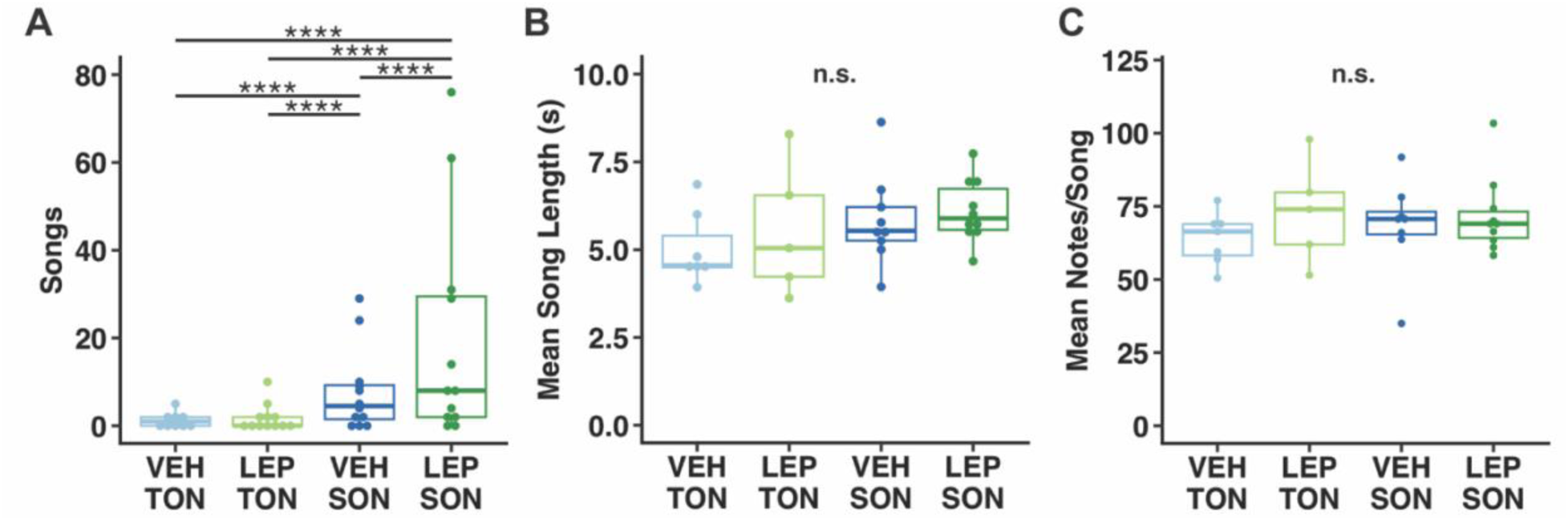
Leptin and song playback influence song effort. **A)** Effect of leptin and song playback on singing rate. Animals receiving song playback significantly increased their rate of singing. Mice that received leptin injection prior to song playback sang significantly more than mice that received vehicle injection. Neither **B)** mean song length nor **C)** mean note number per song differed by hormone or playback treatment. ****p<0.0001

Although both leptin and song playback influenced the rate of singing, they did not influence the duration of songs that were produced (Fig. 2B,C). Two-way ANOVA did not show a significant effect of hormone condition (F_1,27_ = 0.969, p = 0.33), playback condition (F_1,27_ = 2.317, p = 0.14), or hormone x playback interaction (F_1,27_ = 0.081, p = 0.779) on mean song length. Similarly, mean note number in songs did not differ by hormone condition (F_1,26_ = 1.277, p = 0.27), playback condition (F_1,26_ = 0.104, p = 0.749), or hormone x playback interaction (F_1,26_ = 0.296, p = 0.591). Finally, we found no differences in rate of counter-singing (F_1,21_ = 0.223, p = 0.64) or counter-song latency (t_14.9_ = 0.825, p = 0.42) across hormone treatment among animals that received song playback (Fig. 3).

**Fig. 3.**
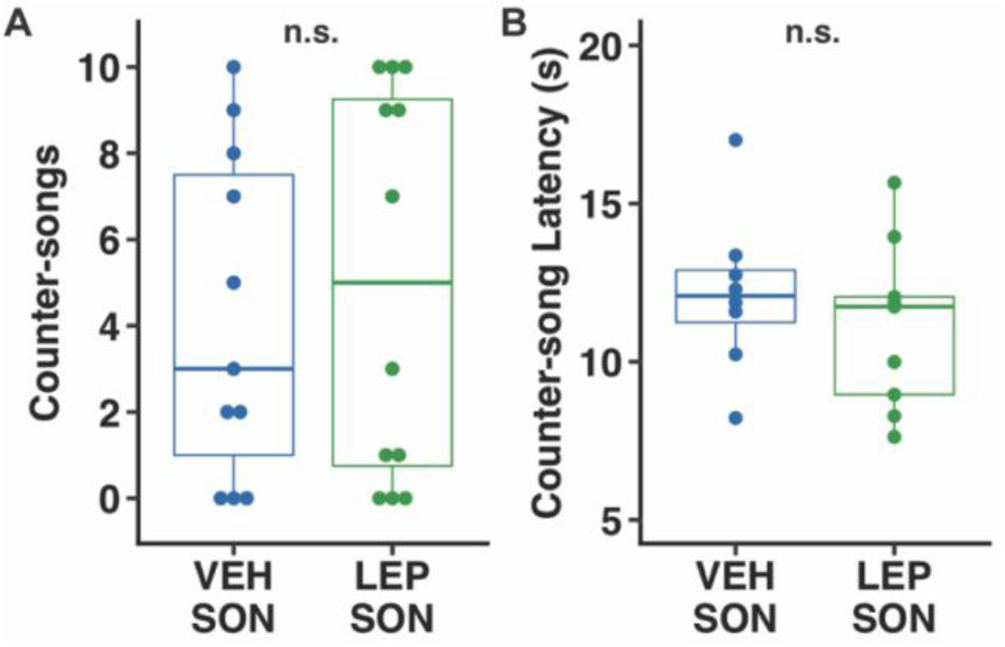
Leptin does not affect counter-singing. Leptin injection did not influence **A)** the likelihood of counter-singing or **B)** the latency to counter-sing following song playback.

### Leptin and song playback influence different aspects of song structure

In addition to measures of effort related to singing rate and song length, we also examined the effects of leptin and song playback on measures of song structure that were previously associated with song effort in wild caught singing mice (Burkhard et al., 2023, 2018). We found that starting inter-note interval was significantly longer in mice receiving song playback compared those receiving tone playback (Fig. 4A). A two-way ANOVA showed a significant effect of playback condition (F_1,26_ = 4.641, p = 0.041), but not hormone condition (F_1,26_ = 0.871, p = 0.36), or hormone x playback interaction (F_1,26_ = 0.002, p = 0.963). *Post hoc* Tukey-tests found a significant difference based on playback type (p = 0.042) but not hormone treatment (p = 0.36), nor any differences in pairwise comparisons between individual groups (p > 0.1 for each comparison).

**Fig. 4.**
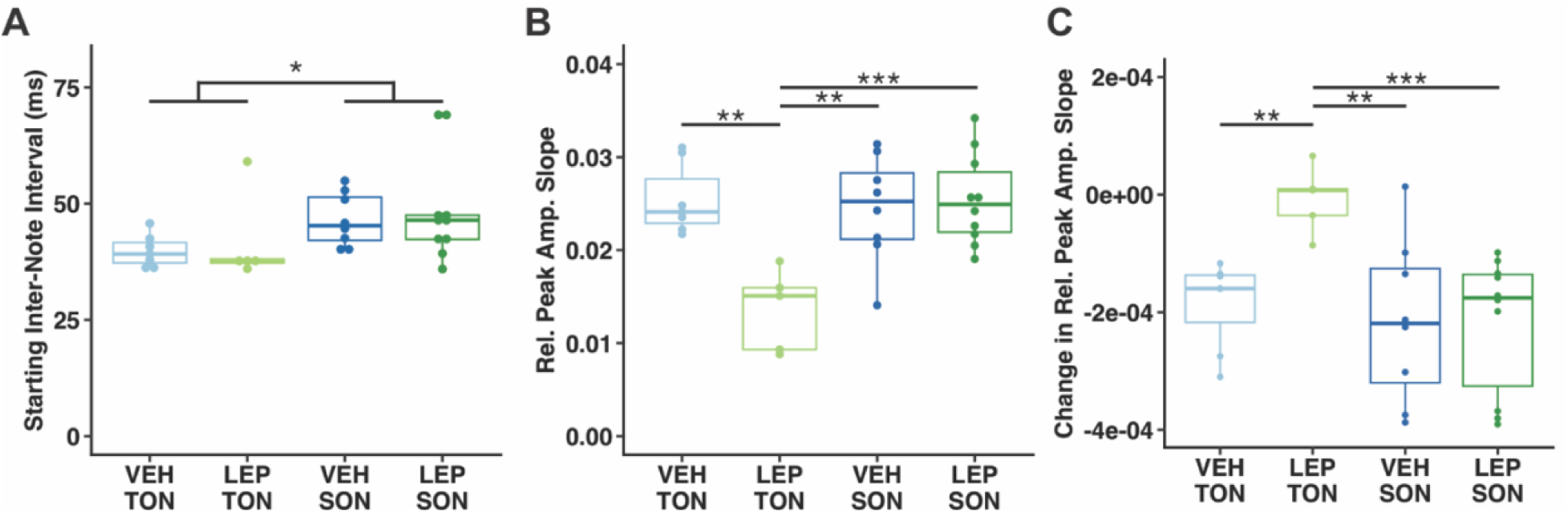
Leptin and song playback influence song structure. **A)** Mice receiving song playback had longer starting inter-note intervals (INIc) than mice receiving tone playback. Mice that receive leptin and tone playback had smaller **B)** starting slope and **C)** rate of slope change for relative peak amplitude compared to all other groups. *p<0.05, **p<0.01, ***p<0.001.

In contrast to other song measures we examined, both measures of amplitude modulation, starting slope (PAb) and rate of change of relative peak amplitude (PAa), were influenced by an interaction between hormone and playback condition (Fig. 4B,C). When testing for effects on PAb, a two-way ANOVA showed a near-significant effect of hormone condition (F_1,26_ = 3.813, p = 0.062) and significant effects of playback condition (F_1,26_ = 7.837, p = 0.0095) and hormone x playback interaction (F_1,26_ = 12.196, p = 0.0017) with *post hoc* tests showing that animals receiving leptin and tone playback differed from all other groups (comparison with leptin and song group, p = 0.00076; for comparison with vehicle and song, p = 0.0028; for comparison with vehicle and tone, p = 0.0016), while there were no significant differences between any other groups (p > 0.95 for all other comparisons). Similarly, when examining PAa, two-way ANOVA found a significant effect of playback condition (F_1,26_ = 8.008, p = 0.0089) and a hormone x playback interaction (F_1,26_ = 4.871, p = 0.036), but no effect of hormone condition alone (F_1,26_ = 1.848, p = 0.19). Once again, *post hoc* tests revealed that animals that received leptin injection and tone playback differed from all other groups (vs. leptin and song group, p = 0.0077; vs. vehicle and song, p = 0.012; vs. vehicle and tone, p = 0.044), with no differences between any other groups (p > 0.90 for all other comparisons).

### Leptin influences non-vocal behaviors

Because leptin signals high energy balance, we also examined its effects on behaviors related to energy expenditure and ingestive behavior. We found that wheel running time, a proxy for increased non-social energy expenditure, did not significantly differ across hormone and playback treatments (Fig. 5A). A two-way ANOVA found no significant effect of playback condition (F_1,44_ = 1.096, p = 0.30) or hormone x playback interaction (F_1,44_ = 0.127, p = 0.72); however, there was a trend toward a significant effect of hormone condition (F_1,44_ = 2.892, p = 0.096), with animals treated with leptin (mean ± standard deviation = 1282 ± 711 s) tending to spend more time running on the wheel than animals treated with vehicle (mean ± standard deviation = 922 ± 744 s).

**Fig. 5.**
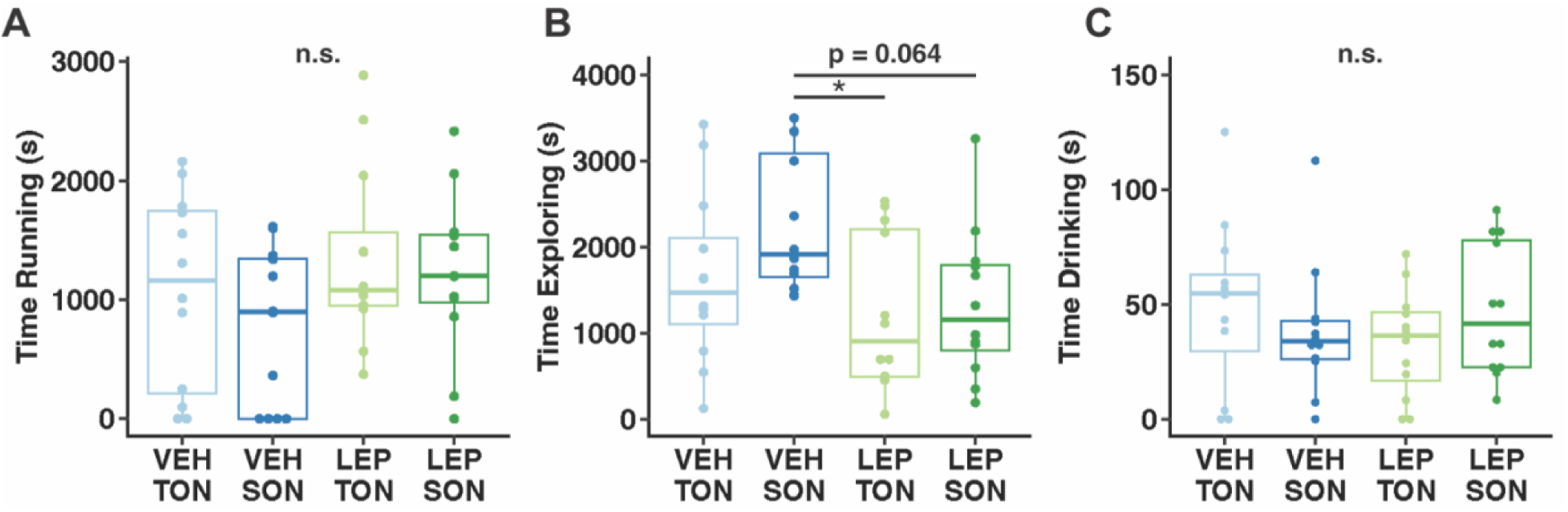
Non-vocal behavioral responses to hormonal and playback manipulations. **A)** Hormone and playback type did not influence time spent running on an exercise wheel. **B)** Mice that received vehicle injection explored the cage significantly more than mice that received leptin injection. **C)** Neither hormone nor playback manipulation affected amount of time spent drinking water. *p<0.05.

Conversely, mice that received vehicle injection spent significantly more time exploring the cage, defined as actively moving around the cage but not running on the wheel, compared to animals that received leptin injection (Fig. 5B). A two-way ANOVA showed a significant effect of hormone treatment (F_1,44_ = 6.777 p = 0.013), while we did not find a significant effect of playback condition (F_1,44_ = 2.026, p = 0.16), or a hormone x playback interaction (F_1,44_ = 1.041, p = 0.31). *Post hoc* tests revealed that mice given vehicle and song playback spent significantly more time exploring than mice given leptin with tone playback (p = 0.033), with a near significant trend for exploring more than mice given leptin with song playback (p = 0.064).

While we observed a significant decrease on exploring time and a trend toward increased wheel running time for leptin-treated mice, we did not find a significant effect of leptin on time spent drinking water (Fig. 5C), with a two-way ANOVA showing no significant effect of hormone treatment (F_1,44 =_ 0.177, p = 0.68), playback treatment (F_1,44_ = 0.039, p = 0.84), or hormone x playback interaction (F_1,44_ = 2.30, p = 0.137).

### Measures of song effort co-vary

Finally, we sought to identify whether any of the measures of song production, features associated with song effort, or non-vocal behaviors consistently co-varied with one another (Fig. 6). We found that the number of songs produced during playback was significantly positively correlated with average song length (rho = 0.63, FDR = 0.0016) as well as number of counter-songs produced (rho = 0.81, FDR = 8.53×10^-7^) and significantly negatively correlated with counter-song latency (rho = −0.72, FDR = 0.0027). We also found that counter-song latency was positively correlated with starting inter-note interval (rho = 0.74, FDR = 0.0023) and identified a strong negative relationship between starting relative peak amplitude slope and relative peak amplitude rate of change (rho = −0.87, FDR = 2.04×10^-8^).

**Fig. 6.**
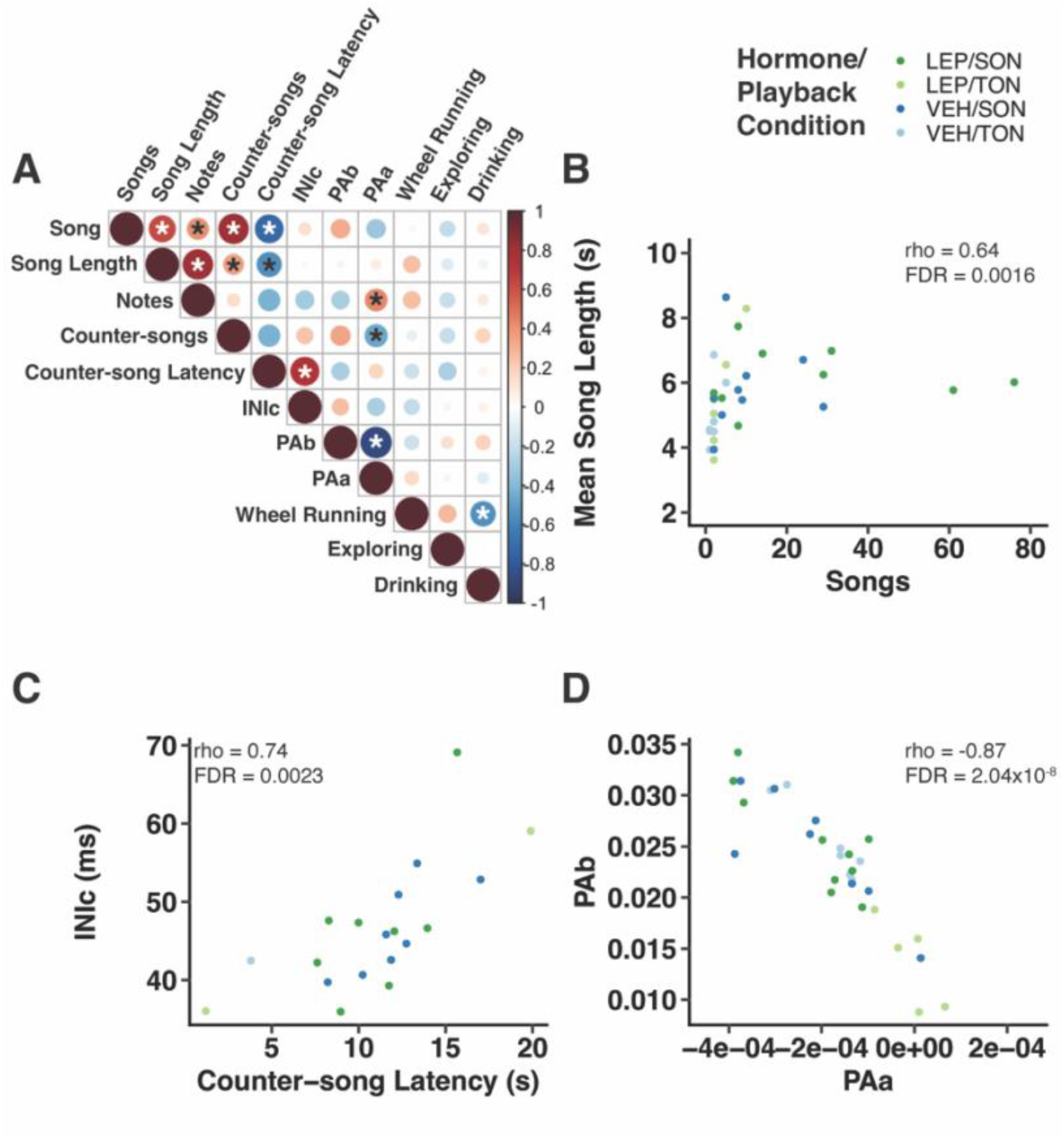
Song features associated with effort co-vary. **A)** Correlations between vocal behavior, song features, and non-vocal behaviors. Color indicates value of correlation coeffection (Spearman’s rho) and circle size indicates its absolute value. Asterisks show significant correlations (p<0.05) and white asterisks indicate correlations that remained significant (FDR<0.05) after correcting for multiple tests. **B)** Relationship between number of songs produced during playback and mean song length. **C)** Relationship between counter-song latency and starting inter-note interval (INIc). **D)** Relationship between starting slope of relative peak amplitude (PAb) and rate of change of relative peak amplitude slope (PAa). Color of points in B-D indicates hormone and playback condition.

## Discussion

In this study, we investigated the role of leptin and social environment in regulating vocal advertisement display effort in Alston’s singing mouse. We found that animals that received playback of conspecific songs sang significantly more songs than individuals that received tone playback. Among these, the mice that had received leptin injection sang more frequently than those that had received vehicle injection. Surprisingly, however, leptin injection did not influence rate of singing in animals that received tone playback. Instead, we saw that mice given leptin and tone playback differed in measures of amplitude modulation compared to mice in other treatment groups, while playback treatment alone influenced the rate of note production at the beginnings of songs. Together, these results demonstrate that both leptin and perceived social environment are important regulators of acoustic display investment that interact to influence advertisement song effort across multiple dimensions. These findings deepen our understanding of the role of leptin in social communication and raise new questions about how individuals integrate signals of their own internal state with cues from the environment to regulate social behaviors.

Our study builds on prior research into the role of leptin in regulating display effort in singing mice. Previously, in wild caught singing mice, circulating leptin levels were found to be the single strongest predictor of song effort among a panel of physical, hormonal, and nutrient measures related to body condition (Burkhard et al., 2018). Additionally, in a laboratory study, leptin injection resulted in increased singing rates (Giglio and Phelps, 2020). Surprisingly, our current results differ from this previous laboratory study in two key ways. First, that earlier experiment found that leptin injection increased singing rate both before and after receiving playback of conspecific male song. Additionally, they found that both leptin injection and song playback resulted in significantly shorter songs compared to saline injection and playback of tones or silence.

These differences are likely due to variation in experimental design. The previous study used mice with *ad lib* food access and a low dose (0.1 mg/kg) of leptin while we used fasted mice and a much higher (10 mg/kg) leptin dose. There are also significant differences in the playback experiments. While the earlier study used a within-subjects design to observe leptin effects both before and after song playback, in the current study we used a 2 x 2 between-subjects design to investigate leptin and playback effects separately. Additionally, we used more song playbacks over a longer time period and positioned the playback speaker further from the subject mouse (Fujishima and Long, 2025), resulting in much higher song rates. This result suggests that the effects of leptin on song production vary by social context and may be dose dependent.

When examining measures of song length, we were surprised that neither hormone nor playback treatment had a significant effect. Previously, leptin was found to decrease song length and leptin-injected animals sang shorter songs following playback compared to their own songs before playback (Giglio and Phelps, 2020). However, most other studies investigating social effects on singing mouse advertisement display have found that song length increases in response to song playback (Tripp and Phelps, 2025, 2024) or live interactions with conspecifics (Giglio, 2020; Okobi et al., 2019). Based on these prior studies, we expected that leptin and playback would each regulate song length. However, we also recognize that due to the fact that we only included song length data from mice that sang during playback and that several mice—including over half of the subjects given tone playback—did not sing, we may lack the necessary sample size and resulting power to detect smaller differences in song length.

In addition to re-examining leptin and playback effects on song rate and length in a new paradigm, we also build on earlier studies by investigating how these hormonal and social signals influence other aspects of song production and structure. First, we focused on animals that received song playback to examine leptin’s effects on counter-songs, the rapid and precisely timed songs produced immediately following the end of another mouse’s song. Counter-songs rely on cortical processing and can be sung with latencies as low as one second after stimulus song offset (Okobi et al., 2019). This behavior is still not well understood, but current evidence suggests that it plays a role in home range defense or mutual avoidance of conspecifics (Fujishima and Long, 2025; Tripp and Phelps, 2025). Given these factors, we expected that counter-singing would be sensitive to signals of energy balance. However, we found no difference in the likelihood or latency to counter-sing across hormone treatments in animals given song playback. This suggests that leptin signaling does not strongly influence the likelihood of a mouse to respond to a conspecific song but does increase the level of effort invested in that response by increasing the number of songs produced following playback.

Along with aspects of song production, we also examined how leptin and playback affect features of song structure, focusing on starting inter-note interval and two measures of amplitude modulation. Prior studies of wild-caught subjects used principal components analyses to identify the major axes of variation in singing mouse song, which were categorized as song effort and frequency modulation (Burkhard et al., 2023, 2018). The measures we chose to focus on for this study were among those that loaded most strongly onto the song effort component, with starting inter-note interval (INIc) and slope of relative peak amplitude (PAb) loading strongly and negatively, and rate of change of relative peak amplitude slope (PAa) loading strongly and positively on this component.

There are also *a priori* reasons to expect these values to be related to song effort. In the case of inter-note interval, faster trills would result in smaller values. In singing mice, trill rate is androgen dependent and females prefer to approach songs with higher trill rates, suggesting it is a high effort, honest signal of quality (Pasch et al., 2011). The fact that animals that received tone playback had smaller initial inter-note intervals, then, was surprising, as we expected song playback to evoke higher effort songs. Instead, our results may reveal a trade-off between rate of song production and trill rate within songs. As mice sing more and more songs in response to playback, they may need to reduce the effort devoted to each individual song in the form of reduced trill rate. However, this possibility is diminished by the lack of significant relationship between song rate and starting inter-note interval (Fig. 6A).

In the case of amplitude modulation, both of our measures suggest that mice that were given leptin followed by tone playback had less variation in amplitude throughout their songs compared to those in other groups, with smaller starting slopes and reduced rate of change in relative peak note amplitude. In other words, the mice given leptin and tone playback started their songs with notes closer to the highest amplitude recorded during their playback session. Because these mice scored higher on a measure positively associated with song effort in wild mice (PAa) and lower on a measure negatively associated with song effort (PAb), we interpret this as mice in the leptin and tone group devoting more effort to song amplitude compared to mice in other groups. This would suggest that leptin increases display effort by promoting songs with higher amplitude throughout in the absence of social stimuli, but when mice hear nearby conspecifics, the effect of leptin shifts to increased song rate. However, because we were only able to use relative measures of amplitude, we cannot rule out the possibility that mice in the leptin and tone group sang quieter, lower effort, songs overall.

Beyond effects on advertisement display, leptin treatment also influenced the physical activities mice engaged in during the playback hour. Specifically, we noticed a significant decrease in cage exploration and a trend toward increased wheel running in mice treated with leptin. While we were somewhat surprised by these results, prior literature shows that leptin’s effects on physical activity can be complex. Leptin injection increases spontaneous physical activity and wheel running in fed animals (Choi et al., 2008; Matheny et al., 2009; Morton et al., 2011). However, fasted animals also show increases in ambulatory activity and wheel running, which are reversed by leptin. This increase in activity has been interpreted as food-seeking behavior (Morton et al., 2011) and as singing mice in the present study were fasted, the increased exploratory behavior observed in vehicle-treated animals may be related to food-seeking.

Unlike previous rodent studies, we do not see a similar pattern for wheel running. Instead, our data suggest that leptin treatment of fasted singing mice may weakly promote voluntary exercise in the form of wheel running, though this trend was not statistically significant. Several studies have demonstrated the rewarding effects of wheel running (Belke and Wagner, 2005; Greenwood et al., 2011; Iversen, 1993; Lett et al., 2000; Meijer and Robbers, 2014) and our results may indicate that, for fasted singing mice, leptin shifts activity away from food seeking and toward rewarding exercise.

While leptin has primarily been studied as a regulator of feeding and body weight, we wanted to determine whether leptin injection influenced ingestion generally in fasted singing mice. We found that there was no difference in time spent drinking between animals that received leptin injection compared to those that received vehicle injection. These results are consistent with earlier studies which found no effects of leptin on water intake in rats (Patel and Ebenezer, 2008) or chickens (Denbow et al., 2000), unless they had been selectively bred for high body weight (Kuo et al., 2005).

A major caveat to our findings regarding non-vocal behaviors is that we measured time engaged in each of these activities rather than actual energy expended, distance traveled, or water ingested. Thus, these relatively gross measures may obscure some differences, and these results should be interpreted with caution. Still, the effects of leptin physical activity in singing mice and other wild-derived rodents are worthy of further study.

We also sought to understand how our measures of song production, song effort, and non-vocal behavior related to one another, finding several measures that were significantly correlated. In particular, the number of songs produced in response to playback was significantly associated with mean song length, number of counter-songs, and shorter counter-song latencies. The correlation between song rate and length was somewhat surprising, given the lack of significant differences between treatment groups in song length. However, a closer look at the data shows that animals with very high song rates had roughly average song lengths, suggesting that while mice typically demonstrate high song effort through both increased rate and length of songs, at very high singing rates it may not be possible to continue to sing long songs.

Along with singing longer songs, mice that sang more overall were also more likely to counter-sing and had shorter counter-song latencies. This suggests that the likelihood and latency to counter-sing are indeed indicators of increased investment in song effort, despite the fact that we did not find differences in these measures across hormonal treatment groups. A study with a larger sample size focused on leptin’s effects on counter-singing may reveal a more subtle difference, relative to the change in song rate we found here. Additionally, we observed that mice with shorter counter-song latencies also had shorter starting inter-note intervals, suggesting that mice that sang quickly in response to playback also did so with higher-effort songs.

Finally, we found a negative relationship between starting slope of relative peak amplitude (PAb) and rate of change of that slope (PAa), demonstrating that songs with relatively steep increases in relative amplitude at the beginning of the song are also characterized by faster rates of change between relative note amplitudes. This observation is consistent with our interpretation that mice in the leptin/tone group started their songs closer to the peak song amplitude, resulting in smaller differences in relative amplitude at the beginning of the song and slower rate of change between relative note amplitudes.

In addition to singing mice, the role of leptin in regulating social signaling has also been investigated in weakly electric fish that use electric organ discharges (EODs) in both foraging and social communication. In the glass knifefish (*Eigenmannia virescens*), EOD amplitude, but not frequency, was increased by leptin injection in food deprived animals (Sinnett and Markham, 2015). The fact that leptin most clearly increases song rate in singing mice and EOD amplitude in ghost knifefish may reflect which signal features are most costly in each species. For singing mice, the choice to sing can result in both conspecific and heterospecific aggression (Fujishima and Long, 2025; Pasch et al., 2013) as well as predation (Pasch and Pino, 2013), suggesting that song rate likely has the highest potential cost. On the other hand, *E. virescens* produce EODs at high (approximately 200-600 Hz), individual-specific, rates (Hopkins, 1974), which suggests that changes in frequency come with very little cost or benefit, while increases in amplitude are likely more costly. However, because our recording methods did not allow us to reliably measure absolute song amplitude, which is impacted by the animal’s distance and position relative to the microphone, it remains a possibility that singing mice also modulate their song amplitude in response to leptin signaling.

In addition to social display, leptin has also been found to regulate investment in reproductive behaviors. In male rats, intracerebroventricular administration of leptin results in preference for sexual behavior over caloric intake and measured by an increase in ejaculations and decrease in sucrose solution consumption (Ammar et al., 2000). Similarly, in red-sided garter snakes (*Thamnophis sirtalis parietalis*) intraperitoneal injection of leptin increased both appetitive and consummatory sexual behavior in both sexes during the mating season, in which the snakes do not eat (Wilson et al., 2021). Conversely, in Syrian hamsters leptin decreased lordosis in fasted females but increased lordosis in females that had been fed *ad libitum (Wade et al., 1997)*, suggesting that leptin’s effects may also be species and context dependent.

Along with leptin, there is a striking overlap in the neurohormones and circuits involved in regulating feeding and social behaviors (Fischer and O’Connell, 2017). For example, the neuropeptide galanin promotes feeding (Crawley, 1999), while galanin-expressing neurons of the preoptic area play important roles in mating behavior and parental care (Bakker et al., 2002; Fischer et al., 2019; Kohl et al., 2018; Tripp et al., 2020; Wu et al., 2014). Additionally, oxytocin (and its homologues in non-mammalian taxa), is well known as a regulator of social behaviors including parental care (Dan et al., 2024; O’Connell et al., 2012; Olazábal, 2018), pair bonding (Insel and Hulihan, 1995; Klatt and Goodson, 2013), and vocalization (Goodson and Bass, 2000), but also acts as an appetite suppressant (Leslie et al., 2018). Perhaps most relevant to the current study are the effects of ghrelin, which is released by the stomach and promotes feeding through opposing action to leptin on hypothalamic circuits that regulate energy balance (Date et al., 2000; Wren et al., 2001). In addition to its effect on feeding, ghrelin has been found to suppress ultrasonic courtship vocalizations in mice (Shah and Nyby, 2010), revealing that ghrelin and leptin can play opposing roles in regulating both feeding and social behaviors.

Overall, this study demonstrates that singing mouse display effort depends both on social context and perceived energy balance, signaled by leptin. These results add to a growing literature seeking to understand the connections and interactions between neurohormonal signals of energy balance and social behavior. While there appears to be a particularly strong connection between circuits and hormonal signals that regulate feeding and social behavior, these behaviors are likely also influenced by further environmental cues and signals of internal state. Future studies should further investigate the interactions between environmental and state cues as well as identify the site of action of these signals and how neural circuits regulating feeding and social behaviors interact with one another.

## Acknowledgments

We thank the University of Texas at Austin Animal Resources Center staff for assistance with animal care. This work was supported by NIH BRAIN initiative (M190270174 to S.M.P.) and the National Institute for Mental Health (F32MH125562 to J.A.T.).

